# DoliClock: A Lipid-Based Aging Clock Reveals Accelerated Aging in Neurological Disorders

**DOI:** 10.1101/2024.02.01.578331

**Authors:** Djakim Latumalea, Maximilian Unfried, Diogo Barardo, Jan Gruber, Brian K. Kennedy

## Abstract

Aging is a multifaceted process influenced by intrinsic and extrinsic factors, with lipid alterations playing a critical role in brain aging and neurological disorders. This study introduces DoliClock, a lipid-based biological aging clock designed to predict the age of the prefrontal cortex using post-mortem lipidomic data. Significant age acceleration was observed in samples with autism, schizophrenia, and Down syndrome, with autism showing the most pronounced effects in aging-rate. An increase in entropy around age 40, suggests dysregulation of the mevalonate pathway and dolichol accumulation. Dolichol, a lipid integral to N-glycosylation and intracellular transport, emerged as a potential aging biomarker, with specific variants such as dolichol-19 and dolichol-20 showing unique age-related associations. These findings suggest that lipidomics can provide valuable insights into the molecular mechanisms of brain aging and neurological disorders. By linking dolichol levels and entropy changes to aging, this study highlights the potential of lipid-based biomarkers for understanding and predicting biological age, especially in conditions associated with premature aging.

## Introduction

Neurological syndromes including Parkinson’s and Alzheimer’s disproportionately affect the older population [1]. Moreover, these disorders (and others) are major contributors to mortality [2] and cause a significant financial burden [3]. The primary risk factor for developing most neurological disorders is aging [4]. While aging is characterized by a loss of physiological function and an exponential increase in mortality [5], aging is also a highly heterogeneous process [6] influenced by a range of intrinsic and extrinsic factors [7].

Lopez-Otin et al. [8] proposed a set of hallmarks of aging, which can be further classified into three distinct categories: primary, antagonistic, and integrative hallmarks. These hallmarks aim to capture the underlying causes of damage, the key responses to such damage, and the resulting effects responsible for the ultimate functional decline of aging organisms.

Alterations in lipid composition play a critical role as they are involved in metabolic energy, homeostasis, and cell signaling [9,10]. In addition, aging is known to significantly alter the brain lipidome [11], which will impact the hallmarks of aging, such as proteostasis [12], and result in decreased resilience and neuronal plasticity [13]. It therefore stands to reason that age-dependent alterations in the lipidome may contribute to brain aging and increase the risk of developing neurological disorders [4]. Evidence already exists that lipid alterations contribute to specific age-dependent neurological disorders. For instance, dysregulated lipid homeostasis has been linked to inflammaging which plays a role in the etiology of Alzheimer’s disease [14]. In addition, lipid peroxidation is positively correlated with Alzheimer’s disease, Down syndrome, and other neurological disorders [15]. Although neurological disorders are mostly found in aged individuals, they can develop at a younger age under certain contexts. Prenatal stress and other environmental stressors can increase the risk of developing neurological disorders, such as schizophrenia, and autism spectrum disorders [16–18]. People with Down syndrome, schizophrenia, and autism also tend to die at a younger age [19–21], which may indicate accelerated aging.

Age acceleration can be measured using aging clocks, which may provide insight into the underlying factors that contribute to the aging process. Numerous aging clocks have been developed to predict biological age. The first prominent clock, developed by Hannum et al. [22], was based on DNA methylation data from whole blood. Subsequently, various other aging clocks using DNA methylation data have been developed and proposed [23,24]. This has been followed by the development of aging clocks utilizing other omics data, such as transcriptomics [25], proteomics [26], lipidomics [27], or a combination of various data sources [28,29]. Most of these models are considered first-generation aging clocks as they predict chronological age. However, chronological age may not always be the most accurate marker, which has prompted researchers to directly predict mortality [30,31]. These clocks are known as second-generation clocks.

Horvath et al. [32] also used their aging clock [23] on brain tissue from individuals with Down syndrome showing a significant age acceleration effect. In addition, Cole et al. [33] utilized neuroimaging data to identify factors associated with age acceleration in individuals with Down Syndrome. Both studies found significant age acceleration and showed that age acceleration is measurable through DNA methylation and neuroimaging.

Age acceleration has also been proposed in schizophrenia, although studies are conflicting [34]. Higgins Chen et al. [35] explored 14 epigenetic clocks and found that 3 mortality-based clocks were able to identify significant acceleration in schizophrenia patients. Conversely, other studies utilizing epigenetic clocks based on chronological age or mortality data did not observe significantly accelerated tissue specific brain aging [36,37]. This may be caused by the relatively low number of samples. It seems that DNA methylation clocks lack the capability to measure aging in individuals with schizophrenia, whereas mortality clocks are proficient, although they depend on clinical parameters. Developing an aging clock based on lipids provides another perspective.

Additionally, there is very limited data on brain aging in individuals with Autism [22,38], highlighting a critical gap in our understanding of how Autism may influence the aging process. In this study, we aimed to determine if lipidomics data from the prefrontal cortex could predict biological age. While several clocks have been devised for human brain tissue [32,39,40], they predominantly rely on DNA methylation data, or on transcriptomic data [41,42]. However, to our knowledge, no model based on lipids from the prefrontal cortex has been developed thus far, making this study a novel exploration in the field of aging clocks. Lipids are integral to understanding the relationship between aging and neurological disorders, given that they constitute approximately 40% of the dry-weight gray matter [43], with the brain exhibiting the highest diversity of lipid species [44]. These findings suggest that a lipid-based clock might pinpoint aging-associated lipids, potentially shedding light on conditions like Down syndrome, schizophrenia, and autism. Additionally, we identify molecules strongly linked with age and propose them as predictive biomarkers [45,46]. Our study demonstrates that variations in brain lipids suffice for estimating biological age.

## Results

In the present study, 163 lipid species and 242 samples were retained, selected for unique chemical formulas to ensure dataset granularity. Among 39,446 lipid concentration values, 1.98% were missing, with 47% of lipids having at least one missing value. Missing values were imputed using K-nearest neighbors (k = 5). The dataset included 195 samples without neurological disorder (WND), 27 samples with schizophrenia (SZ), 15 samples with autism spectrum disorder, and 5 samples with Down syndrome (DS). **Supplementary Table 1** provides a comprehensive breakdown of the preprocessed dataset. No outliers were identified.

### Identifying a Robust Model

Twenty-six machine learning models were trained using 100 bootstrap iterations to identify the most generalizable model. Linear regression models demonstrated the best performance (**Supplementary Figure 1**), and detailed model performance metrics are presented in **Supplementary Table 2**). Based on these findings, we developed an Elastic Net model and applied principal component analysis (PCA) to reduce dimensionality and mitigate noise. PCA projections were used as input for the model, a strategy commonly employed in similar studies [27,39,47]. We bootstrapped the data 10,000 times with replacement, stratifying by age group, sex, and ethnicity, and trained Elastic Net models using WND samples. During training, we included ethnicity, sex, and the post-mortem interval (PMI) as covariates to account for additional variance.

### Exploring Principal Components and Lipid Patterns

To investigate whether there was a relationship between the principal components and the metadata, we conducted a correlation analysis. The first principal component showed no substantial correlation with the meta data, but was enriched for PG(0-20:0/22:4), a glycerophospholipid involved in membrane signaling, likely due to its high variance. In contrast, principal components two and three exhibited significant Pearson correlation coefficients (r) with Shannon entropy (r = -0.30, *P*<0.001 and r = 0.30, *P*<0.001, respectively). Dolichol-19 and dolichol-20 were identified as primary contributors to entropy in 31% of samples. When entropy was recalculated using only dolichols, a striking correlation with chronological age (r = 0.92, *P*<0.001) emerged. These findings suggested that dolichols could serve as age-associated markers.

### Entropy and Aging

Samples were divided into six age bins (20-80 years, 10-year interval), and entropy levels were compared using the Wilcoxon Rank Sum test. Samples aged 40-50, 50-60, and 60-70 exhibited significantly higher entropy levels than their younger counterparts (*P* < 0.001 for each group). These results indicate a substantial increase in entropy around the age of 40-50 (**Figure 1A**). Interestingly, no significant differences in entropy were observed between ASD, SZ, DS compared to controls, suggesting that age-related changes in entropy were more pronounced than disorder-specific patterns.

**Figure 1.**
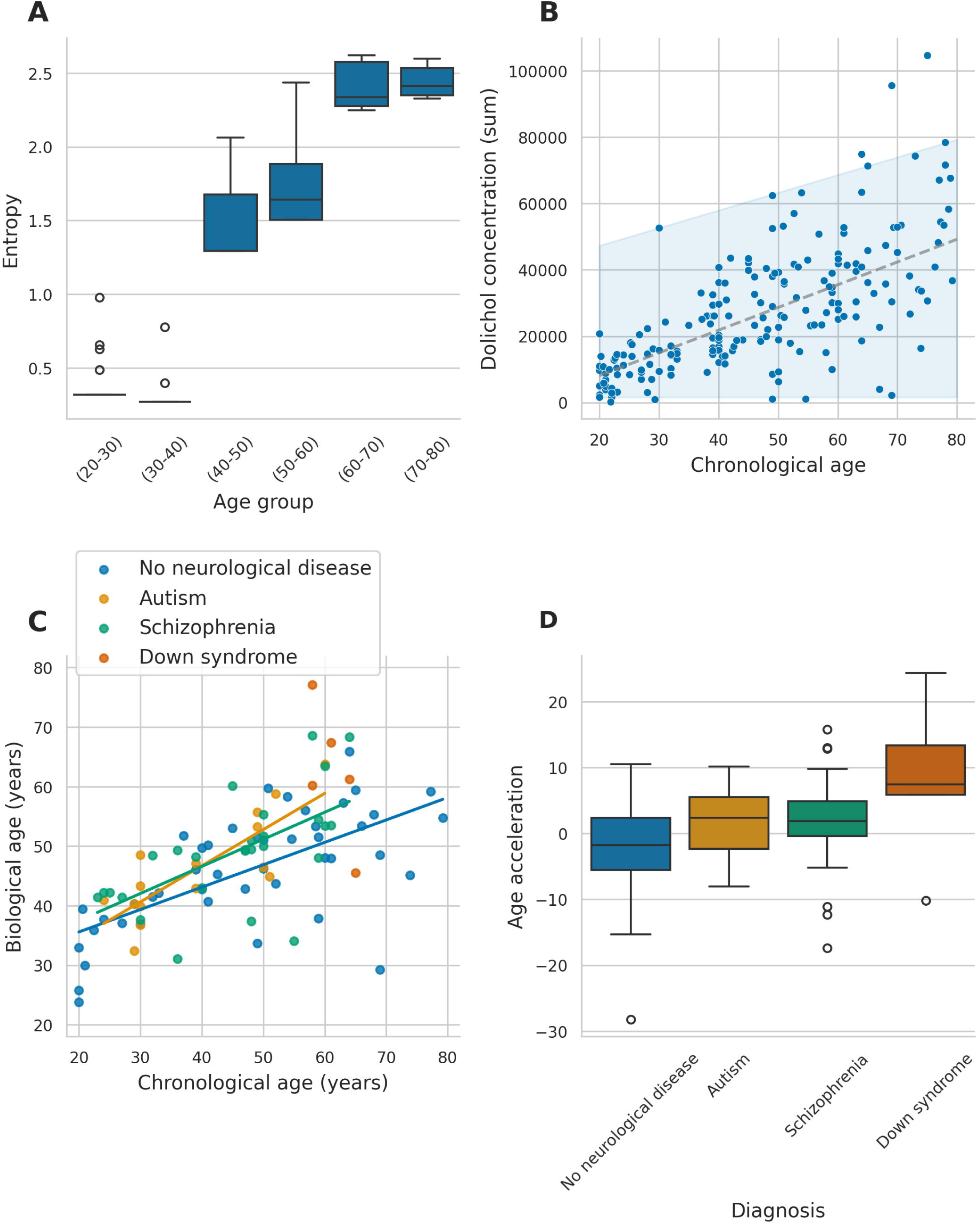
Association between entropy, biological age, and biomarkers across neurological conditions. (A) Boxplots of entropy across different age groups. (B) Behavior of summed dolichol concentration through age, with a 95% confidence interval. (C) Predicted values of the models for all samples, including groups with no neurological disease, autism, schizophrenia, and Down syndrome. (D) Boxplot comparing age acceleration across the different groups.

### Aging-Associated Principal Components

PC5, strongly correlated with age (r = 0.61, *P* < 0.001), stood out as a key feature in the Elastic Net model, receiving the highest coefficient during training. The loadings of PC5 were dominated by dolichol-19, dolichol-20, and specific glycerophospholipids such as PG(17:1(9Z)/0:0) and PG(22:6). These findings indicated that PC5 encapsulates a broader set of aging-related processes beyond dolichol metabolism.

### Ethnicity and Lipid Profiles

Ethnicity also emerged as a significant factor influencing lipid composition. PC6 correlated positively with Han Chinese ethnicity (r = 0.54, *P* < 0.001) and negatively with Caucasian ethnicity (r = -0.32, *P* < 0.001). These results suggest distinct lipid concentration patterns across ethnic groups. In contrast, surprisingly, sex exhibited no significant correlation with any principal component, underscoring its limited role in lipid variance within this dataset.

### Lipid Trends with Age

Furthermore, we conducted pairwise comparison between different age groups using the Mann-Whitney U test to identify molecules significantly associated with age **(Supplementary Table 3**). Initial findings indicated numerous molecules exhibiting significant increases with age after multiple testing correction using the Bonferroni test, particularly between age groups 0-20 years and 20-40 years. Dolichols exhibited notable increases across multiple age groups, showing a linear trend with age and increasing variance, as depicted in **Figure 1B**. Specifically, dolichol-19 C95H160NO and C95H157O, as well as dolichol-20 C100H164ONa, C100H165O, C100H168NO, demonstrated significant increases between age groups 0-20, and 20-40 (*P* < 0.00007 for each), and between groups 20-40 and 40-60 (*P* < 0.00007 for each). However, only dolichol-20 C100H164ONa exhibited a significant increase between groups 40-60 and 60-80 (*P* < 0.00007), with no significant increases observed between groups 60-80 and 80-100, potentially due to limited sample size and high variance. The log fold changes are shown in **Supplementary Table 4**.

Based on these findings, it was hypothesized that dolichol may serve as a biomarker for predicting biological age, given its significant and consistent increase across age groups.

### DoliClock

Building on our findings, we hypothesized that dolichols could serve as reliable biomarkers of age. To test this, we developed DoliClock, an Elastic Net model trained exclusively on dolichol lipids. The model incorporated sex, ethnicity, and PMI as covariates to account for additional sources of variation. Using the default Elastic Net parameters, we performed 10,000 bootstrap iterations stratified by ethnicity and sex. The model achieved a median absolute error of 8.96 years, with the median-performing model selected for further analyses. The distribution of the performance can be found in **Supplementary Figure 2**.

DoliClock was subsequently applied to predict the ages of samples with neurological disorders, as shown in **Figure 1C**. We compared the intercepts and slopes of the trendlines for WND, SZ, and ASD samples using a t-test, excluding DS samples due to their limited size in comparable age ranges. As illustrated in **Figure 1C**, the aging slope for ASD samples (0.61) was significantly steeper than that for WND samples (0.38, *P* = 0.048), indicating a 1.6-times increase in the rate of aging in ASD samples. In contrast, the slope for SZ samples (*P* = 0.28) did not differ significantly from that of WND samples.

To further evaluate aging dynamics, we calculated age acceleration (**Figure 1D**). ASD, SZ, and DS samples exhibited significantly greater age acceleration compared to WND samples (*P* = 0.047, *P* = 0.008, and *P*=0.015, respectively), suggesting these conditions are associated with accelerated aging.

Feature importance analysis using SHAP values identified dolichol-20 C100H164ONa as the most influential predictor of biological age (**Figure 2**). Lower concentrations of dolichol-20 and dolichol-19 were linked to younger predicted ages, while moderate or high concentrations corresponded to older predicted ages. This relationship between dolichol and chronological age changes over time, as can be seen across the different age groups; in younger samples, dolichol levels are associated with a lower biological age, whereas in older samples, dolichol levels are indicative of biological aging due to their increase in concentration levels.

**Figure 2.**
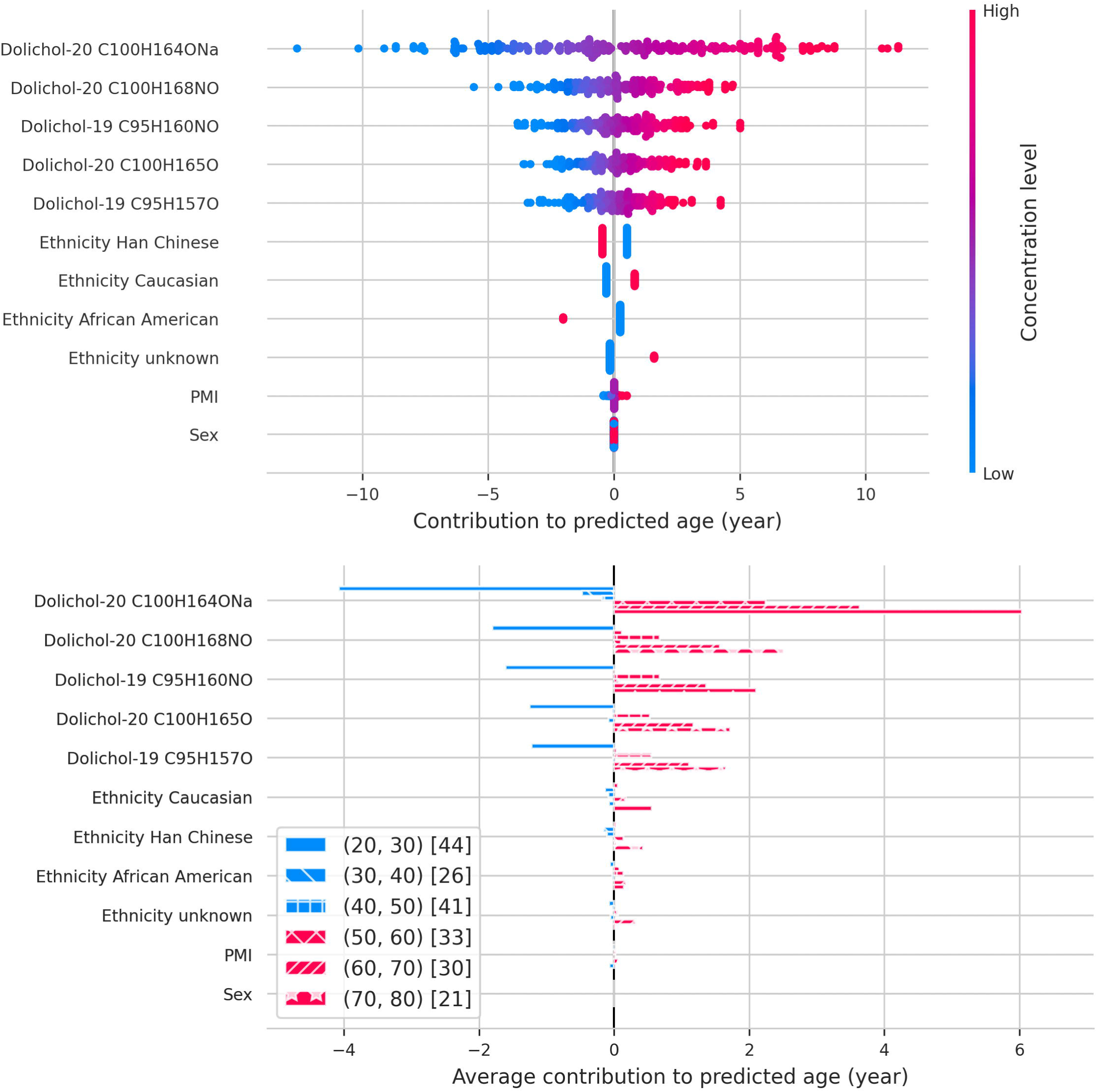
SHAP values showing the contribution of each feature to the predicted age. The plots illustrate the relative importance and direction of influence of each feature on the age predictions.

Ethnicity had a minor but measurable effect on DoliClock predictions. Han Chinese samples were generally predicted to be younger, whereas Caucasian samples were predicted to be older. However, differences in age distributions across ethnic groups in the training data likely contributed to these patterns (**Supplementary Figure 3**).

Finally, an analysis of dolichol variance revealed that variance increases with age (**Figure 3**). This trend was consistent across the lifespan, with a pronounced rise in variance observed around the age of 40 (*P* < 0.001 for all dolichols, Levene’s test). This suggests that age-related changes in dolichol levels not only reflect chronological aging but also increase heterogeneity, potentially offering further insights into individual biological aging trajectories.

**Figure 3.**
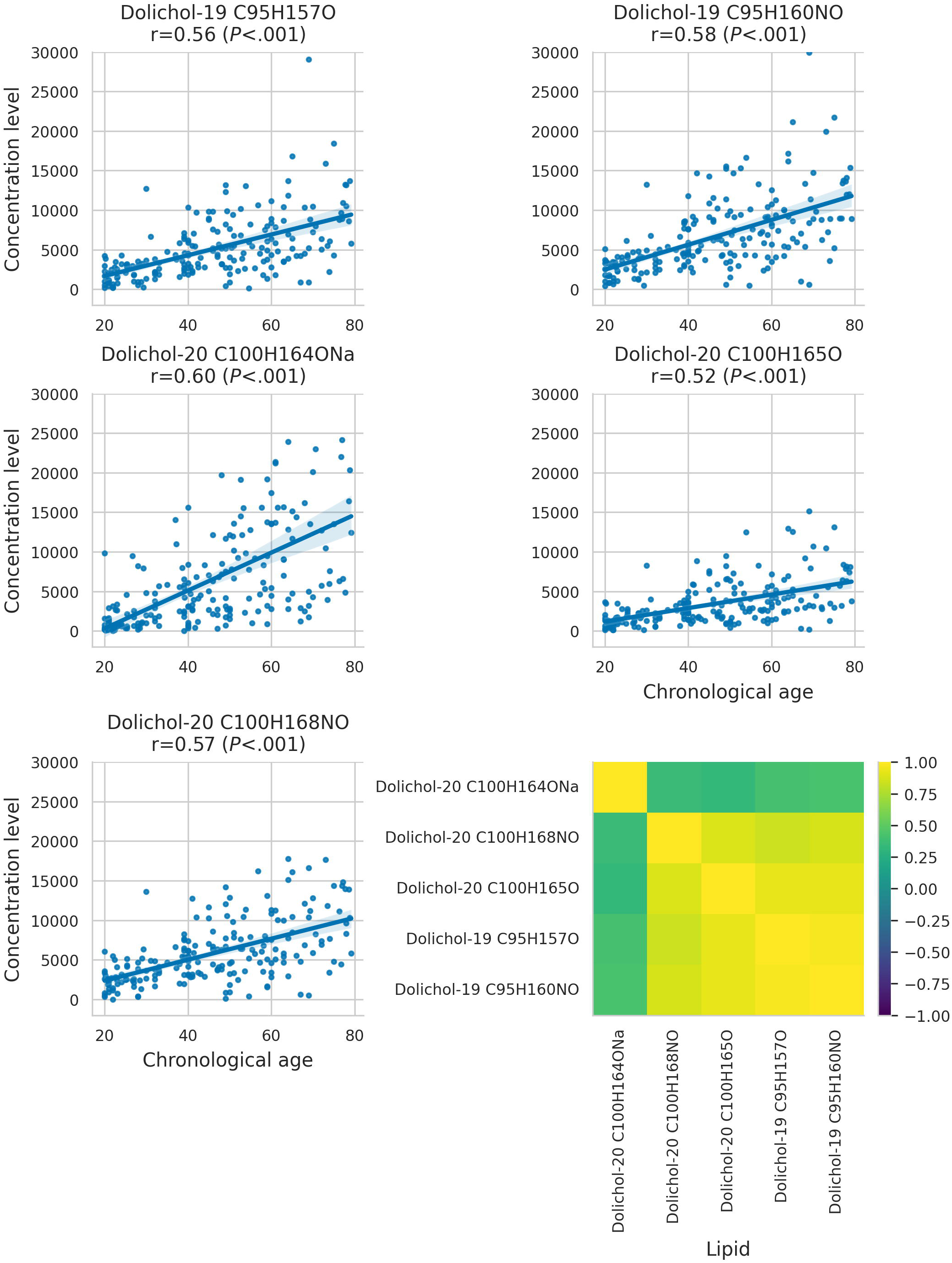
Behavior of dolichol concentrations for each isoprenoid and chemical formulation, shown separately. The figure also highlights the correlations between different isoprenoids and formulations.

## Discussion

### Age acceleration

Our study identified a significantly faster aging rate in samples with autism and significant age acceleration in samples with autism, schizophrenia, and Down syndrome.

The findings for schizophrenia align with and extend conflicting outcomes reported in prior studies using epigenetic clocks, which assess chronological age and mortality [35–37].

Lipidomic profiling, as applied in this study, appears to capture distinct age-related signatures in schizophrenia, offering an alternative perspective on aging processes in this population.

Similarly, while further exploration into brain aging in autism remains necessary [20,38], our results demonstrate significant age acceleration in individuals with autism. This aligns with prior research suggesting a potential role of genetic factors linked to both aging and autistic traits or the indirect influence of these traits on lifestyle, thereby modulating the aging process [48]. For samples with Down syndrome, our findings corroborate earlier studies that documented premature aging through epigenetic markers and neuroimaging [32,33]. This consistency highlights the utility of lipidomic analysis in understanding age-related changes in Down syndrome.

### Entropy and Age-Related Dysregulation

Our study highlights a striking escalation in entropy, particularly around the age of 40. This suggests a possible dysregulation in the mevalonate pathway, potentially leading to the accumulation of dolichol.

The observed strong correlation between entropy and chronological age supports the hypothesis that entropy reflects age-dependent dysregulation in dolichol-related pathways. Dolichols, as members of the polyprenol class, are essential for intracellular transport [49], N-glycosylation [50], and play a pivotal role in the mevalonate pathway [51], which synthesizes key molecules such as cholesterol, dolichol, and ubiquinone. Alterations in the concentration levels of these molecules, particularly the age-related increase in dolichol [52–56], suggest a dysregulation of the mevalonate pathway. This dysregulation may be mediated by an age-related increase in HMG-CoA reductase, the rate-limiting enzyme of this pathway [57]. These findings reinforce the association between entropy and mevalonate pathway dysregulation and underscore the need for deeper exploration of the role of dolichols in aging.

### Dolichol as an Aging Biomarker

Dolichol’s potential as an aging biomarker is supported by extensive research across various organisms, including mice [58–60], rats [52,61–66], drosophila [67], and humans [53–55,68–71]. Dolichol, a compound comprising a variable number of isoprene units depending on the species [72], has garnered attention for its association with aging processes [73,74]. Studies have reported an accumulation of dolichol in multiple tissues with age, although concentrations may vary among tissue types. For instance, research on rats’ liver revealed that dolichol concentration increases in tandem with HMG-CoA reductase levels [62], with caloric restriction shown to retard dolichol accumulation [63,64]. Similarly, investigations on brain tissue have shown a several-fold increase in dolichol concentration during aging and development, suggesting potential developmental roles [52]. Studies in humans have echoed these findings, with dolichol levels increasing dramatically in adults compared to neonates [68]. Notably, specific dolichol molecules, such as those with isoprene unit 17 up to 21, have been found to be particularly abundant in brain tissue [69]. However, studies have also linked elevated dolichol levels to neurodegenerative diseases, suggesting a potential link to lysosomal dysfunction [53]. Additionally, individuals with conditions like Down syndrome and autism have been observed to exhibit elevated dolichol levels [54,55], further supporting its role as a biomarker of both aging and neurological conditions.

Dolichol also plays critical roles in organelle transport and N-glycosylation [49,50], serving as the mevalonate pathway’s end product [51]. Its role as a lipid carrier for glycan precursors suggests a potential link between glycan and dolichol biomarkers, hinting at a mechanistic relationship [75,76]. Additionally, caloric restriction has shown promise in reducing dolichol accumulation, although its effectiveness may vary depending on the duration and timing [62,63]. Emerging evidence also suggests dolichol may protect aging membranes against free radical damage, potentially slowing biological aging [64]. Conversely, an unstable dolichol system could accelerate aging processes [77]. Another hypothesis posits that dolichol accumulation might involve low-density lipoprotein dysregulation, impairing lysosomal function leading to autophagic degradation [64,78].

### Tissue-Specific Versus Systemic Aging

While our study utilized brain tissue, the regression on chronological age raises questions about whether observed changes reflect tissue-specific aging or systemic aging processes. Brain tissue might uniquely capture certain age-related signatures, but further research is needed to determine whether similar patterns occur in other tissues. Distinguishing between tissue-specific and systemic effects is crucial for understanding the role of dolichol in aging.

### Limitations and Confounding Variables

The post-mortem nature of our dataset necessitates consideration of confounding factors. Cause of death, medication usage, and post-mortem interval (PMI) could all influence lipid levels. Notably, the disproportionately high number of samples with a PMI of exactly 14 among healthy individuals suggests this value might be a default placeholder, potentially introducing bias. Additionally, the unknown medication status of individuals with schizophrenia, a condition often treated with lipid-altering drugs, could further confound results [79]. Addressing these factors in future research will be critical for refining dolichol’s utility as an aging biomarker.

### Future directions

Future research should prioritize validating the findings of this study through external datasets or experimental approaches, given the inherent limitations of post-mortem tissues.

Understanding the mechanisms linking dolichol and aging, particularly the functional distinctions between dolichol-19 and dolichol-20, is crucial for unraveling their roles in the aging process.

Investigating their potential contributions to autophagic degradation and lipid peroxidation may also clarify their involvement in cellular aging.

Additionally, a key avenue for exploration is the comprehensive analysis of dolichol levels across various tissues to uncover clinical aging markers or surrogate indicators for brain aging. Since dolichol concentrations vary significantly between tissue types, integrating lipidomic data from multiple sources could enhance the precision and generalizability of age-related biomarkers.

Furthermore, the interplay between entropy, dolichol, and aging presents an exciting frontier. The observed entropy increase and dolichol-19 concentration shift around age 40 suggest that combining insights from information theory and lipidomics could yield novel perspectives. This interdisciplinary approach may uncover unique molecular signatures of aging, offering deeper insights into its underlying biological mechanisms.

### Conclusion

This study explored the potential of lipidomic data to predict biological age in the prefrontal cortex and to identify critical lipid molecules linked to aging. We introduced DoliClock, a novel tool leveraging prefrontal cortex lipid profiles to estimate age. Notably, DoliClock uncovered previously unreported isotopic and chemical variations in dolichol molecules associated with age, highlighting its capacity to detect subtle, age-related biological effects.

DoliClock demonstrated robust performance in estimating biological age across a spectrum of individuals, including those with neurological disorders like Down syndrome, schizophrenia, and autism, which are commonly associated with accelerated aging phenotypes. Our findings revealed significant age acceleration in autism, schizophrenia, and Down syndrome, with autism cases showing a particularly pronounced increase in the rate of aging.

Furthermore, we identified a notable increase in entropy around age 40, with individuals with neurological disorders exhibiting elevated entropy levels at younger ages. These results emphasize the potential of lipidomic data to inform biological age prediction and underscore its utility in advancing our understanding of lipid-based biomarkers of aging.

## Materials and Methods

### Samples

This study utilized the initial dataset by Yu et al. [11], comprising 452 samples from various brain banks: NICHD Brain and Tissue Bank, Maryland Psychiatric Research Center, Maryland Brain Collection Center, Netherlands Brain Bank, Chinese Brain Bank Center, Harvard Brain Tissue Resource Center, and Autism Tissue Program. Samples, primarily gray matter from the anterior prefrontal cortex, weighed approximately 12.55 mg (±1.65). The dataset included 403 samples without neurological disorders (WND), 5 with Down syndrome (DS), 17 with autism spectrum disorder (ASD), and 27 with schizophrenia (SZ). Age ranges varied: WND samples from 0 to 99 years (median age 24 years, excluding 12 prenatal samples), DS samples from 58 to 65 years (median age 61 years), ASD samples from 18 to 60 years (median age 30 years), and SZ samples from 23 to 64 years (median age 48 years). Post-mortem intervals (PMI) ranged from 0 to 44 hours, with median values of 14, 6.2, 20.3, and 19.5 hours for WND, DS, ASD, and SZ samples, respectively. Ethnicity data were incomplete, labeled as unknown, thus limiting its use for matching case and control groups. Detailed sample statistics are provided in **Supplementary Table 5.** A comprehensive description of these data and the methodology can be found in the publication of Yu et al. in 2018 [11].

### Data analysis

#### Data preprocessing

To standardize ages, we adjusted reported values by subtracting a gestational period of 0.767 years. Subjects younger than 20 clustered in feature space (**Supplementary Figure 4**), likely reflecting ongoing brain development, while those over 80 were mostly Caucasian. To address these biases, we limited our analysis to individuals aged 20 to 80. This filtering yielded 5,024 lipid species, of which 2,222 were annotated with LIPID MAP IDs (LM IDs). To manage degenerate LM IDs, we computed a conformity index, resulting in 360 unique lipid species mapped to LM IDs, molecular weights, and retention times. Isotopic variations were reviewed for discrepancies in m/z values, and mass searches against LIPID MAPS adduct lists were conducted across lipid classes in both ionization modes. Any unmatched values were treated as missing, resulting in a final dataset with 242 samples and 163 features.

Outliers were identified by running PCA and calculating the pairwise Euclidean distances between samples. The distances were then averaged per sample. An outlier was defined as any distance below the first percentile - 3 * the interquartile range or above the third percentile + 3 * the interquartile range.

#### Dimensionality reduction, mutual information, and interpretation

We applied principal component analysis (PCA) for dimensionality reduction, using singular value decomposition (SVD) to decompose the data’s correlation matrix into eigenvalues and eigenvectors, represented as □ = □□□^□^ [80]. PCA was implemented via the scikit-learn library [81] for visualization and subsequent analysis. To further explore relationships among features, eigenvectors were rescaled based on mutual information [82], calculated following the method of Platt et al. [83]. Elastic net was employed to identify sparse, informative features while minimizing overfitting [84]. Using the ElasticNet class from scikit-learn [81], PCA loadings were multiplied by elastic net coefficients to determine feature importance.

#### Bootstrapping, Entropy, and Statistical tests

To ensure robust model evaluation, we applied bootstrapping with stratification on age (5-year bins), sex, ethnicity, maintaining demographic diversity across resampled datasets. Shannon entropy [85], was calculated to assess information content, under the assumption of stable prefrontal cortex function among those aged 20-40 (**Supplementary Figure 5**). Lipid data were log-transformed using the Yeo-Johnson PowerTransformer [81,86], with values coded as 0 or 1 depending whether they are within two standard deviations from the mean. Probabilities were computed within 10-year windows, and entropy for each sample was calculated accordingly. Statistical analysis included the Wilcoxon Rank Sum Test for comparing entropy and lipid levels between age groups, with Bonferroni correction applied (*P* < .00007) for multiple tests.

Trendlines were generated using ordinary least squares, and slopes were compared via T-test. Variance was compared using Levene’s test.

### Implementation Details

The analysis pipeline was implemented in Python (v3.9.7, (59)). Numpy (v1.25.2, [87]), Pandas (v2.1.1, [88]) and Scipy (v1.11.3, [89]) were used for computational purposes and scikit-learn (v1.5.2, [81]) was used for the development of machine learning models. For visualization matplotlib (v3.8.0, [90]) and seaborn (v0.13.2, [91]) were used. For interpretation of models the SHAP library (v0.43.0, [92]) was used.

Age acceleration was calculated by fitting a linear regression model to the predicted age and actual age, then obtaining the residuals. Code to reproduce the results are made available at https://github.com/ddlatumalea/DoliClock-2.0.

## Abbreviations

WND: without neurological disorder
SZ: schizophrenia
ASD: autism spectrum disorder
DS: Down syndrome
PCA: Principal Component Analysis
r: Pearson correlation coefficients
PMI: post-mortem interval

## Author Contributions

Djakim Latumalea: Conceptualization, Methodology, Software, Validation, Formal analysis, Investigation, Data Curation, Writing – Original Draft, Visualization.

Maximilian Unfried: Conceptualization, Methodology, Writing - Review & Editing Diogo Barardo: Conceptualization, Methodology, Writing - Review & Editing Jan Gruber: Conceptualization, Methodology, Writing - Review & Editing

Brian K. Kennedy: Conceptualization, Methodology, Writing - Review & Editing

## Table legend

**ASupplementary Table 1.** Lipids and demographics of the processed dataset, including sample counts by neurological disorder status (No Neurological Disorder, Down Syndrome, Autism, Schizophrenia). Gender, age groups, ethnicity, and postmortem interval (PMI; median ± SD assumed to be in hours) are detailed. Dashes (“-”) indicate unavailable data.

**Supplementary Table 2.** Model performance metrics for all 26 models, including R², RMSE, MAE, explained variance, and MedAE, calculated for both training and test datasets.

**Supplementary Table 3.** P-values from Mann-Whitney U tests comparing groups of 20 years, corrected using a Bonferroni adjustment with a significance threshold of 7.67 × 10┚□.

**Supplementary Table 4.** Log fold changes of dolichol concentration across different groups.

**Supplementary Table 5.** Demographics of the original dataset, including sample counts by neurological disorder status (No Neurological Disorder, Down Syndrome, Autism, Schizophrenia). Gender, age groups, ethnicity, and postmortem interval (PMI; median ± SD assumed to be in hours) are detailed. Dashes (“-”) indicate unavailable data.

## Figure legend

**Supplementary Figure 1.** Comparison of 26 models based on their R² test score versus training score. Each point represents a model, illustrating its performance on both the training and test datasets.

**Supplementary Figure 2.** Distribution of all model scores, with the median value indicated.

**Supplementary Figure 3.** Histogram showing the age distribution across different ethnicities. Age is divided into bins of 10 years, and each ethnicity is represented by a distinct color to highlight demographic variation within the dataset.

**Supplementary Figure 4.** Scatter plots of dimensionality reduction techniques applied to the dataset: (A) PCA, (B) PCA with Mutual Information (MI) rescaling, (C) t-SNE, and (D) UMAP. Each plot visualizes sample clustering based on all lipids, with axes representing the first two dimensions of the respective method.

**Supplementary Figure 5.** Distributions of dolichol concentration over time. Each plot highlights upper and lower limits defining the healthy range, with samples outside two standard deviations marked in green.

