## Supplementary figures and images for "DoliClock: A Lipid-Based Aging Clock Reveals Accelerated Aging in Neurological Disorders"

### Supplementary Figure 1

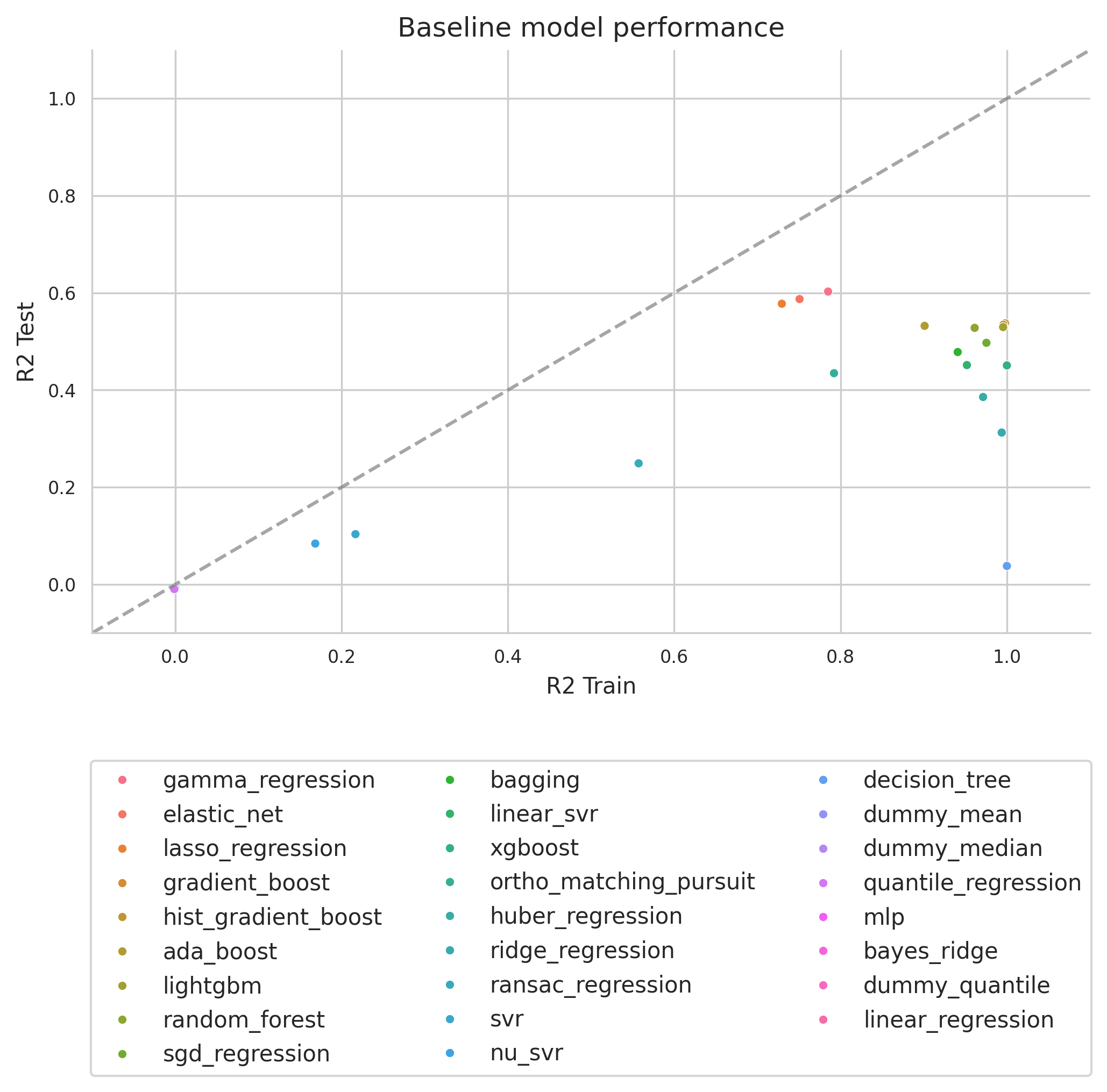

### Supplementary Figure 2

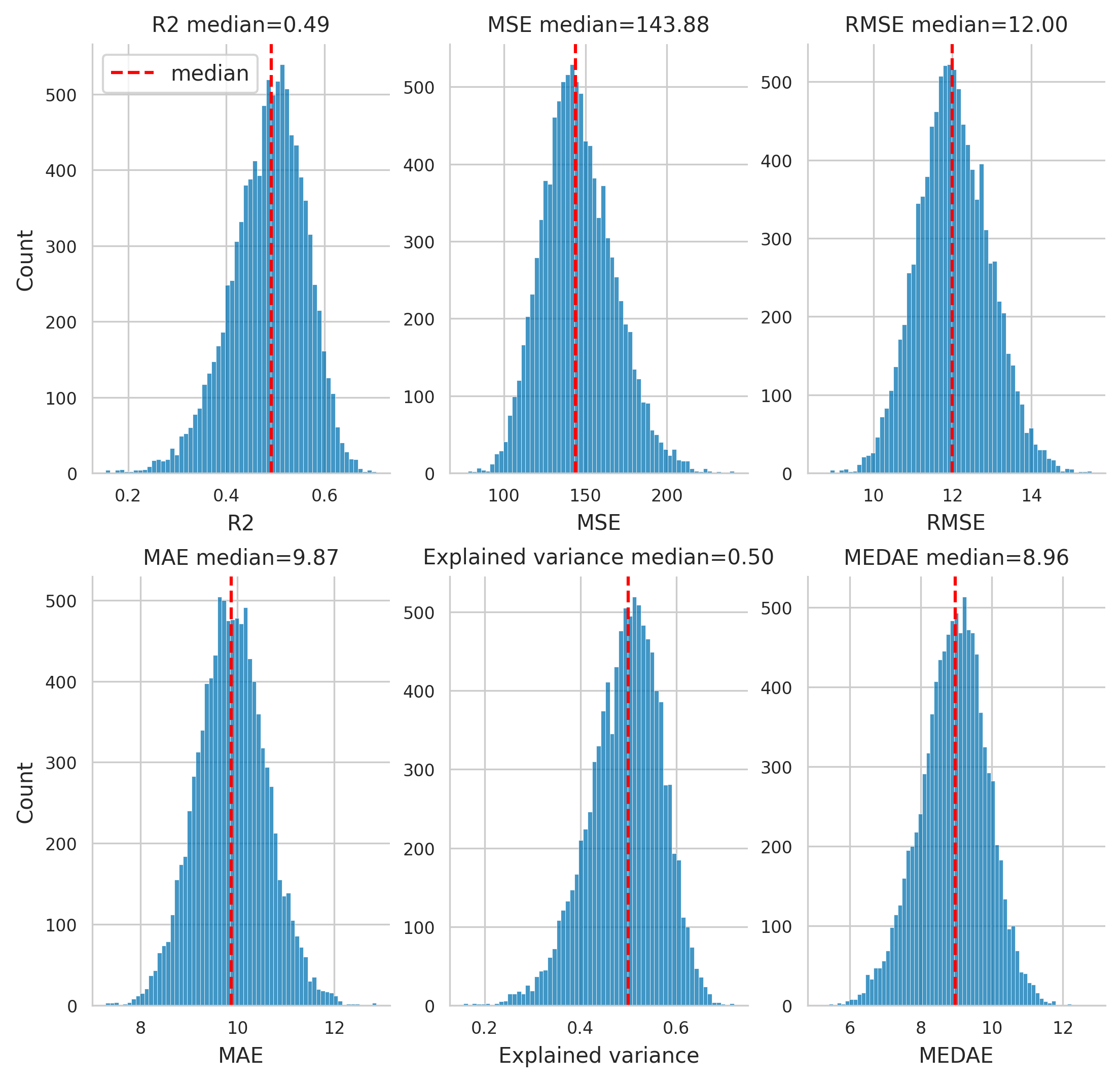

### Supplementary Figure 3

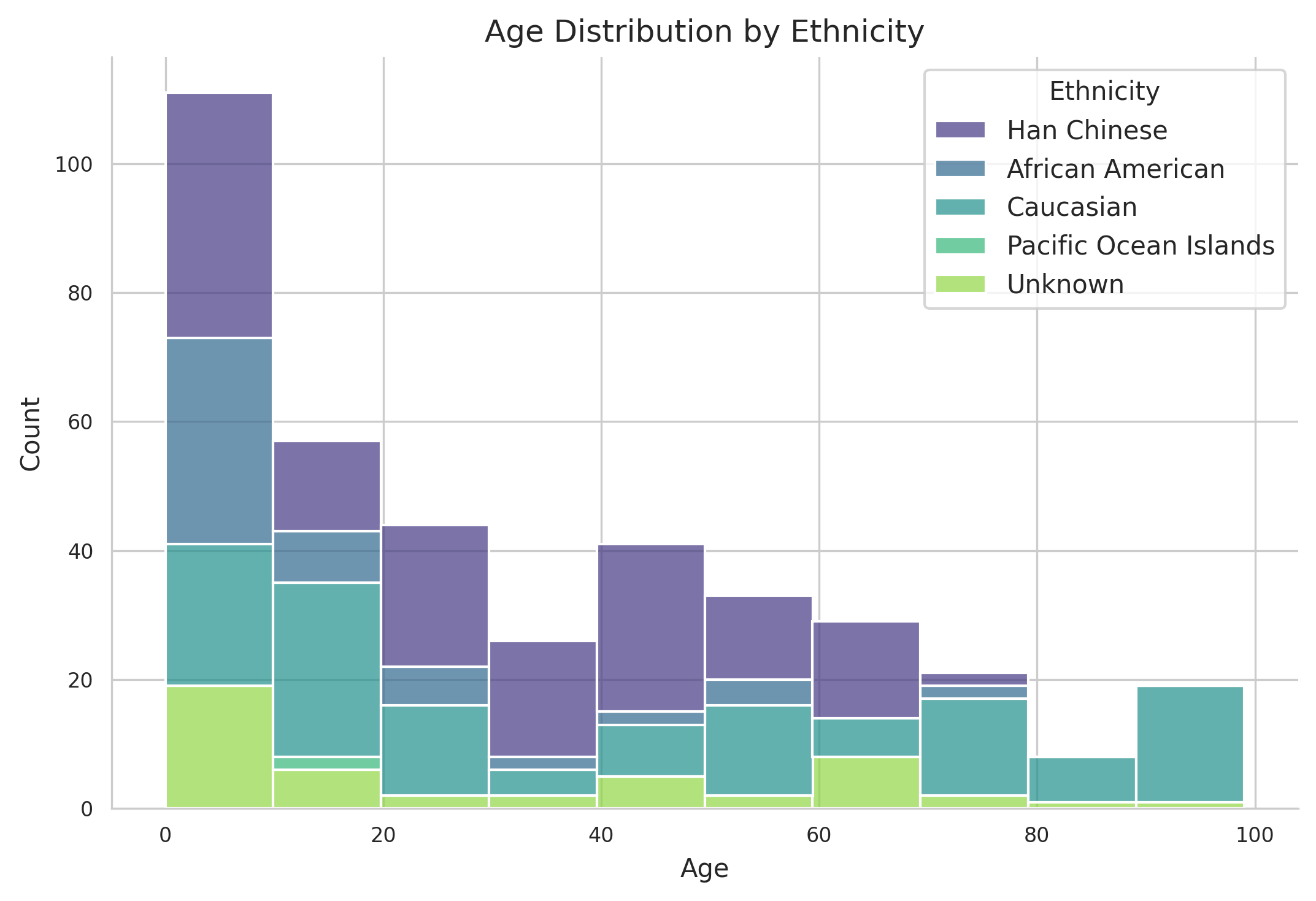

### Supplementary Figure 4

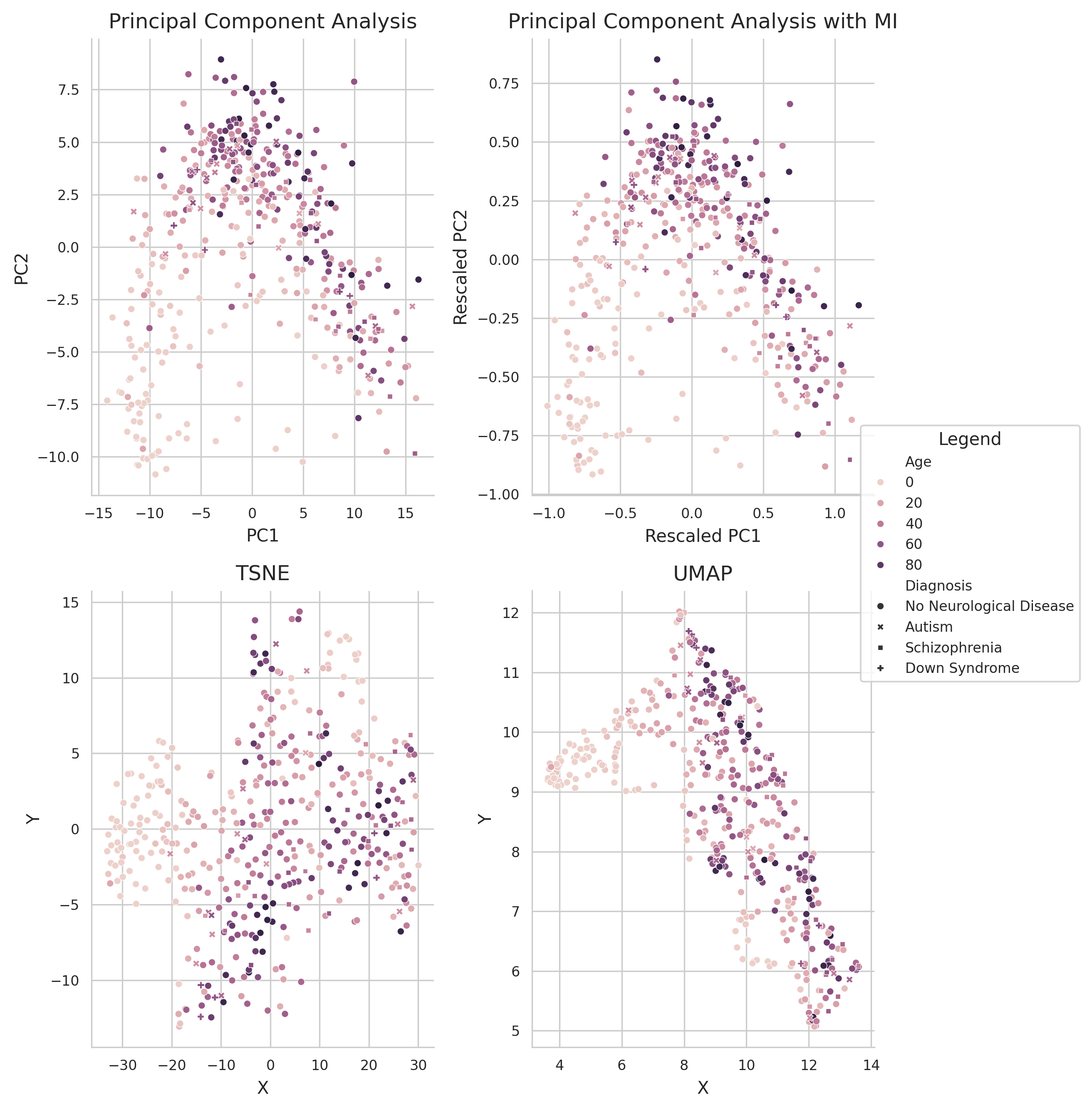

### Supplementary Figure 5

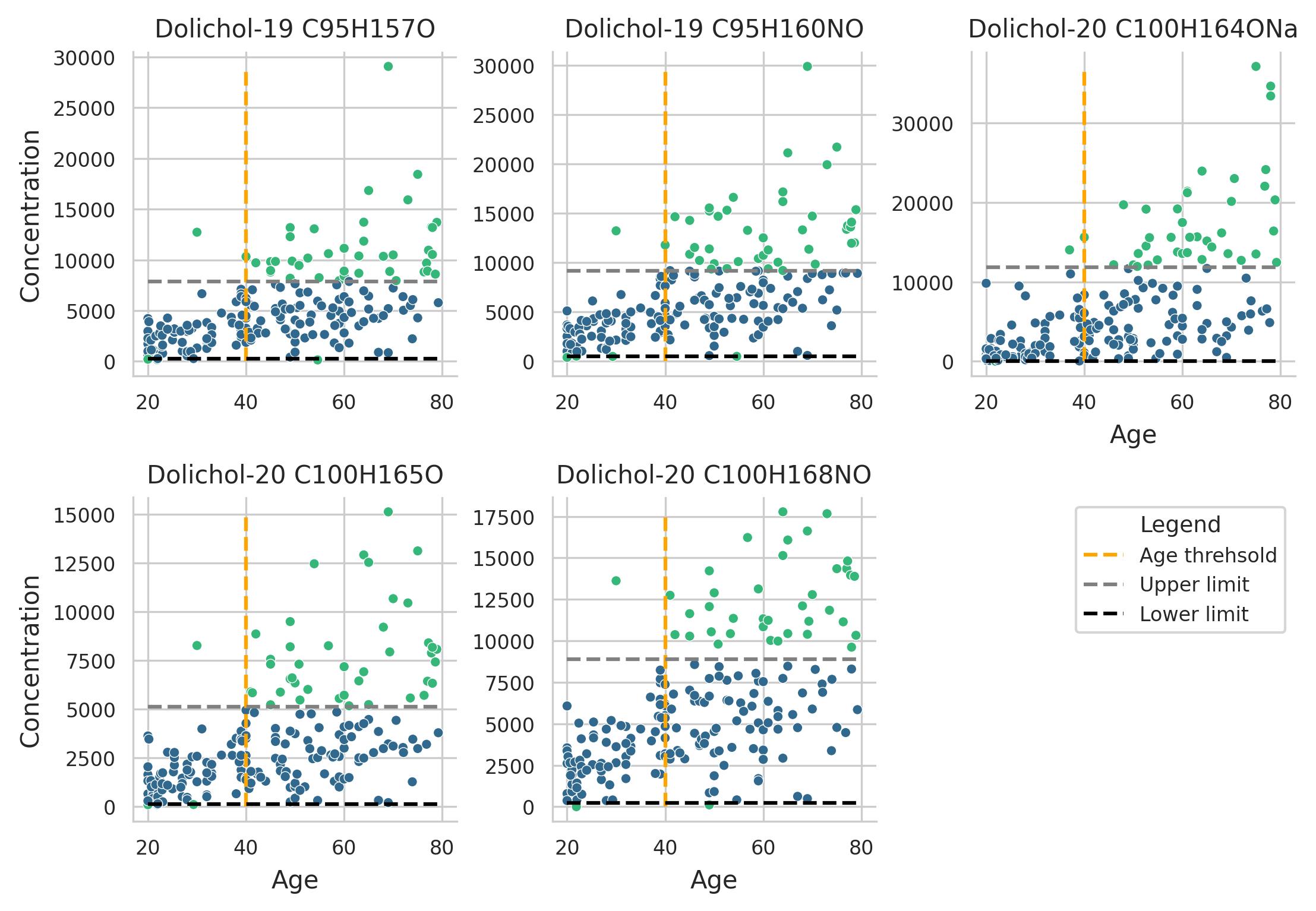
